# Vascular ultrasound for *in vivo* assessment of arterial pathologies in a murine model of atherosclerosis and aortic aneurysm

**DOI:** 10.1101/2022.12.22.521641

**Authors:** Alexander Hof, Henning Guthoff, Maysam Ahdab, Max Landerer, Jasper Schäkel, Jana Niehues, Max Schorscher, Holger Winkels, Simon Geißen, Matti Adam, Martin Mollenhauer, Dennis Mehrkens

**Author notes:** Authors contributed equally (Alexander Hof, Henning Guthoff). **Corresponding Author:** Alexander Hof, MD.

## Abstract

**Background:** Vascular diseases like atherosclerosis or aortic aneurysms are common pathologies in the western world, promoting various, potentially fatal conditions. Hence, a plethora of animal models have been developed to investigate underlying mechanisms and potential therapeutics. Here we evaluate high resolution (HR) ultrasound in mouse models of atherosclerosis and abdominal aortic aneurysm (AAA) for noninvasive monitoring of morphological and functional vascular changes *in vivo*.

**Methods:** Eight-week-old *ApoE*^-/-^ mice were used for disease models. For induction of atherosclerosis, mice were fed a western diet over 12 weeks. To trigger AAA development, osmotic minipumps were implanted, permanently releasing Angiotensin II continuously for 28 days. All animals were on C57Bl6/J background. HR vascular ultrasound of the carotid artery or the abdominal aorta was performed, respectively. Images obtained were analyzed by a speckle tracking algorithm (VevoVasc software) and were correlated with histological analyses by Picro Sirius Red staining and automated collagen quantification.

**Results:** Arterial wall distensibility and global radial strain (GRS) as measures of arterial wall elasticity were reduced in the carotids of atherosclerotic mice as well as in the aortas of AAA mice. Pulse wave velocity (PWV) was elevated in both disease models. Intima-media thickness (IMT) was significantly increased in the atherosclerosis model. Matching those findings, area of the tunica media was enlarged in *ApoE*^-/-^ mice fed a western diet, and in Angiotensin II treated mice as measured by automated image analysis, depicting higher collagen depositions in diseased arteries. Simple regression analysis revealed a strong correlation of media collagen content and area in AAA with IMT and GRS, respectively. In atherosclerosis, media collagen content significantly correlated with PWV and GRS, whereas wall distensibility was associated with the size of media area.

**Conclusion:** Vascular imaging using latest generation HR ultrasound devices is suitable to trace changes of arterial wall properties in murine models of atherosclerosis and AAA. Obtained results not only correlate with histological findings but deliver information on functional parameters which may be used as early disease and risk markers in a longitudinal experimental approach.

## Introduction

Atherosclerosis and aortic aneurysms are severe pathologies of the vasculature, both of which limit prognosis by propagating potentially fatal conditions such as aortic dissection or myocardial infarction [1] [2]. Hence, investigation and revelation of underlying pathomechanisms is of highest relevance.

Several animal models have been established to mimic and investigate mechanisms of cardiovascular disease development [3] [4]. Although histological assessments of arterial disease phenotypes have long been established, postmortem analysis only allows for evaluation of vasculature under static conditions and sample preparation may cause artifacts. In addition, hemodynamic factors like blood pressure, blood flow or viscosity and fluctuating concentrations of various endocrine mediators may influence vascular function *in vivo*, which elude histological examination [5] [6]. Hence, *in vivo* analysis of the vasculature is indispensable.

Vascular ultrasound has emerged as a diagnostic option frequently used for *in vivo* assessment of vascular function in animal models. An obvious advantage over postmortem analysis is the evaluation of multiple vascular parameters in a longitudinal approach, applicable, for example, in the context of drug testing or evaluation of dose tolerance and efficacy at different timepoints in the same animal. In addition to plain determination of vascular wall morphology, physiological parameters like pulse wave velocity (PWV) or arterial elasticity can be assessed. Over the past years, technical features of ultrasound devices have drastically improved, allowing accurate vascular measurements even in small animal models. The Vevo3100 system was specifically designed for preclinical studies and animal models, providing an improved image quality by automated respiratory gating and high resolution (HR) imaging. By using ultra high frequencies with a maximum of 70 MHz, a spatial resolution of up to 30 μm can be achieved with this system, thus enabling the investigator to visualize and trace extremely small structures within the vessel wall [7].

Here, we aim to evaluate and validate the sonographic assessment by Vevo3100 in a murine model of two of the most relevant vascular pathologies, atherosclerosis and aortic aneurysm. The results obtained are compared to histological analyses and automated collagen quantification as described previously [8].

## Methods

### Animal models

Eight-week-old male Apolipoprotein E knock out (*ApoE*^-/-^) mice on C57Bl6/J background were fed a western diet (sniff, 1.25% cholesterol diet, E-15723-347, Soest, Germany) for 12 weeks to induce atherosclerosis **(Fig 1A I)**. Generation of *ApoE*^-/-^ mice was described previously [9]. For induction of abdominal aortic aneurysm (AAA), *ApoE*^-/-^ mice were treated with angiotensin II (Ang II) for 28 days *via* osmotic minipumps as outlined below in detail. C57Bl6/J wildtype (WT) animals served as controls. All animal studies were approved by the local Animal Care and Use Committees (Ministry for Environment, Agriculture, Conservation and Consumer Protection of the State of North Rhine-Westphalia: State Agency for Nature, Environment and Consumer Protection (LANUV), NRW, Germany) and conformed to the guidelines from Directive 2010/63/EU of the European Parliament on the protection of animals used for scientific purposes (81-02.04.2018.A216 and 81-02.04.2020.A249).

**Figure 1:**
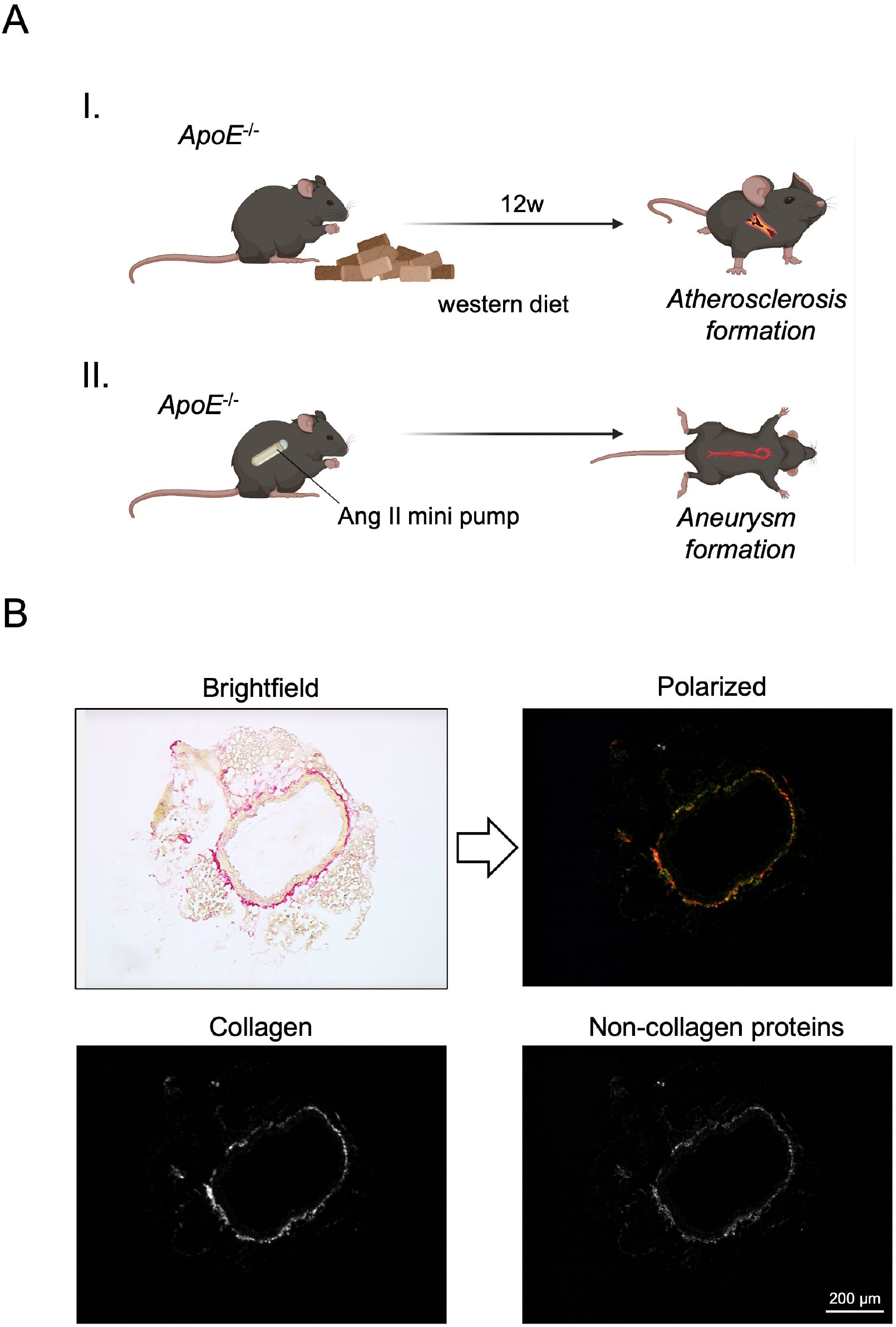
Mouse models and histological workup. *ApoE*^-/-^ mice were fed a high cholesterol western diet for 12 weeks for induction of atherosclerosis (A I). Alternatively, Angiotensin II (Ang II) was infused *via* osmotic minipumps for 28 days to provoke formation of abdominal aortic aneurysm in *ApoE*^-/-^ mice (A II). Aortic sections of the aorta were stained with Picro Sirius Red and visualized in brightfield and polarized light for collagen quantification as described before [8]. Binarization by splitting red and green channels was utilized to separate collagen (red) and non-collagen proteins (green). White pixels in the binarized images were counted to determine total collagen within the samples.

### Implantation of osmotic minipumps

For induction of AAA formation, osmotic minipumps were implanted in eight-week-old *ApoE*^-/-^ mice **(Fig 1A II)**. Alzet osmotic minipumps (Model 2004; ALZA Scientific Products, Cupertino, California, USA) were filled with Ang II (Sigma Chemical Co., St. Louis, Missouri, USA) that subcutaneously delivered 1,000 ng/min/kg of Ang II for 28 days. Mice were anesthetized by isoflurane inhalation (Isofluran-Piramal®, Piramal Critical Care, Voorschoten, The Netherlands; 5% vol/vol for induction and 2% vol/vol for maintenance of anaesthesia) and analgetic treatment *via* subcutaneous injection of buprenorphine (TEMGESIC®, Indivior Europe Limited, Dublin, Ireland; 0.1 mg per kg body weight) prior to implantation. Subsequently, minipumps were implanted subcutaneously into the left flank of the mice.

### Ultrasound analysis

After 12 weeks of western diet feeding, mice were anesthetized with 5% isoflurane and kept under anesthesia with 2% isoflurane. Ultrasound analysis was performed under continuous monitoring of vital signs. Sequences of the carotid artery were acquired reaching from the aorta to carotid bifurcation in B-Mode, M-Mode and ECG-triggered kilohertz visualization (EKV) using a Vevo3100 device and an MX550S transducer (VisualSonics, Fujifilm, Tokio, Japan). For examination of the abdominal aorta after 28 days of Ang II infusion, mice were fixed in prone position and sequences of the longitudinal axis of the abdominal aorta were acquired from paravertebral in B-Mode, M-Mode and EKV. B-Mode images of the carotid artery and the abdominal aorta were examined using VevoVasc software (VisualSonics, Fujifilm, Tokio, Japan). Respiratory gating was performed to minimize artifacts, subsequently borders of the vessel wall and intima-media layer were defined. Strain analysis by advanced speckle tracking of moving image sequences allowed evaluation of elastic properties, in particular wall distensibility and global radial strain (GRS), of the vessel wall. Further, PWV as a measure of arterial stiffness was defined by analyzing ECG-triggered excursion of the arterial wall at two distinct locations of a pre-defined distance in longitudinal acquisitions. Examiners were blinded to genotypes and treatments during acquisition and analysis.

### Tissue preparation

Abdominal cavity was opened under deep anesthesia and analgesia as described above and animals were euthanized by cardiac exsanguination from the left ventricular cavity and consecutive perfusion with ice-cold phosphate buffered saline (PBS). Consecutively, aortas were carefully excised and fixed in 3.7% formaldehyde solution for two days and subsequently embedded in paraffin for histological analyses. Serial sections of tissue specimen of 6 μm thickness were prepared and mounted on microscope slides.

### Picro Sirius Red staining

Slices were thawed and heated to 60°C in a heating chamber for 30 minutes. After rehydrating the samples, Picro Sirius Red (PSR) staining was performed using the Picro Sirius Red Stain Kit (ab150681, Abcam, Cambridge, UK) according to the manufacturer’s instructions. In brief, samples were completely covered with PSR solution in a humid chamber for 60 minutes, rinsed twice in acetic acid and were finally dehydrated using ethanol.

### Collagen quantification

Total collagen content of PSR stained sections was quantified by an elaborated analysis protocol using MatLab as described before [8]. PSR stained sections were imaged by a Leica microscope (Leica DM 4000B, Leica Biosystems Inc., USA) and a polarized filter (Leica ICT/Pol, Leica Biosystems Inc; **Fig. 1B**). Congruent bright-field and polarized images of aortic sections were converted to grayscales by MatLab and media as well as adventitia were automatically defined. White pixels in the binarized, polarized image were counted and their percentage of all pixels classified as media or adventitia was calculated to determine total collagen within these anatomic areas.

### Statistical analysis

Data analysis was performed with student’s t-test and simple regression analysis using GraphPad Prism 8.4.0 (GraphPad Software, San Diego, CA, USA). Data are presented as mean ± SEM. *P*-values < 0.05 were assumed significant. * *p* < 0.05; ** *p* < 0.01; *** *p* < 0.001.

## Results

### Morphologic and functional alterations of the carotid artery in atherosclerosis manifest in vascular ultrasound measurements

To assess properties of the arterial wall in *ApoE*^-/-^ mice after 12 weeks of western diet, ultrasound analysis of the carotid artery was performed as described above **(Fig. 2A-C)**. Intima-media thickness (IMT) was numerically increased in atherosclerotic *ApoE*^-/-^ mice compared to controls (91.4 ±1.3 μm vs. 100.4 ±7.6 μm; **Fig. 2D**). In addition, PWV was significantly higher in those animals (0.98 ±0.03 m/s vs. 1.5 ±0.07 m/s; **Fig. 2E**), suggesting arterial stiffening as expression of the atherosclerotic phenotype.

**Figure 2:**
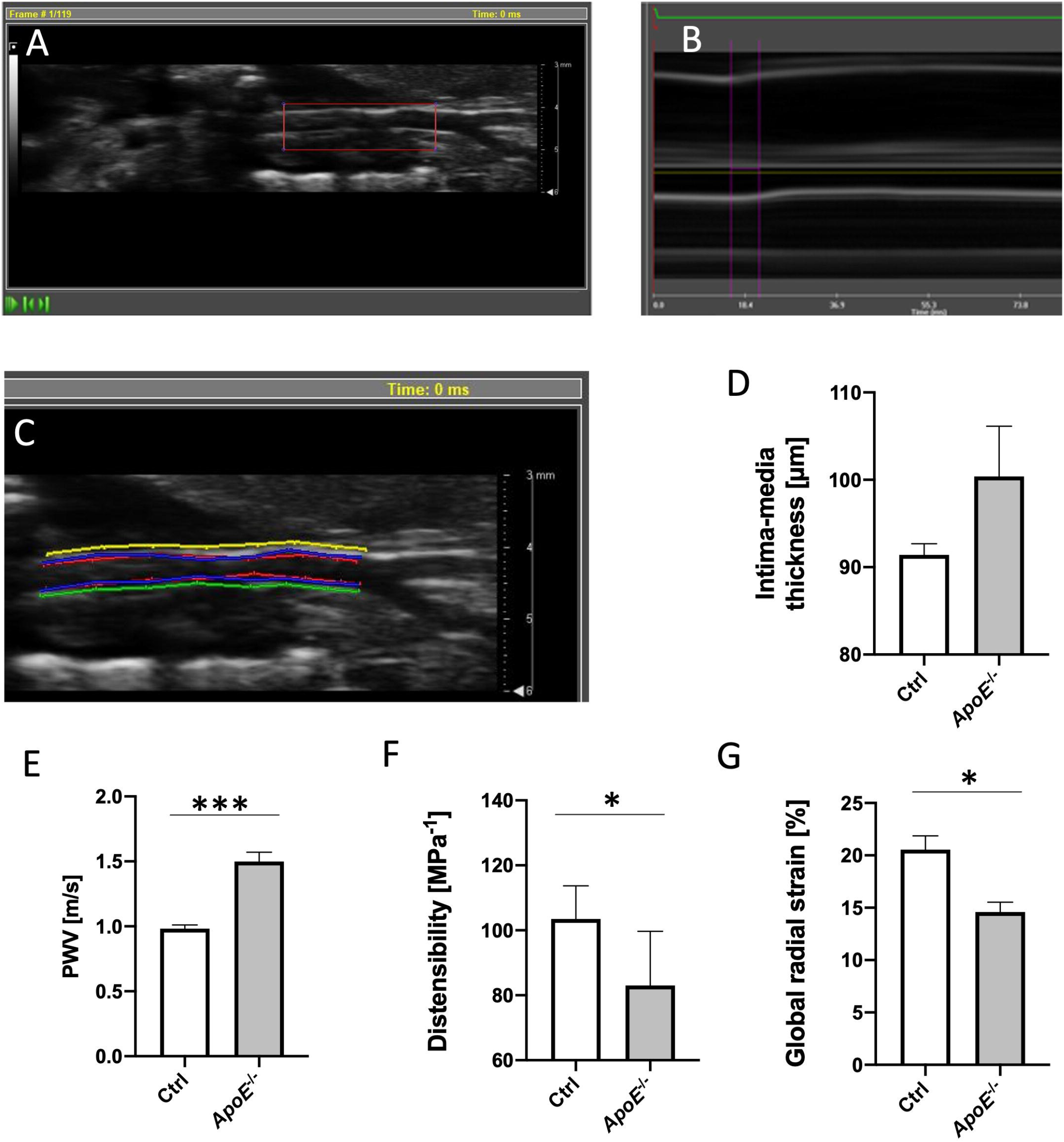
Evaluation of vascular morphology and functional state in atherosclerosis by ultrasound. Segments of the carotid artery were examined from its origin to carotid bifurcation (A, longitudinal acquisition of carotid artery). Pulse wave velocity (PWV) was examined in M-mode, measuring time differences in vessel excursion in different segments of the carotid artery (B). Vessel demarcation was performed at luminal and adventitial site and distensibility and strain analysis was performed by speckle tracking using VevoVasc software (C). Intima-media thickness (IMT) was numerically enlarged in western diet fed *ApoE*^-/-^ mice compared to C57Bl6/J control animals (D). PWV was significantly elevated in carotid arteries of atherosclerotic mice (E), while distensibility (F) and radial strain (G) was significantly lower in *ApoE*^-/-^ mice, indicating reduced elasticity of the vascular wall. *n*=3-4; *p<0.05; ***p<0.001.

Likewise, parameters of arterial wall elasticity were altered in *ApoE*^-/-^ mice fed a western diet. Distensibility of the arterial wall was significantly reduced (103.5 ±10.2 MPa^-1^ vs. 83.0 ±16.7 MPa^-1^) and GRS was diminished (20.6 ±1.3% vs. 14.6 ±0.9%) in atherosclerotic mice **(Fig. 2F, G)**.

### Vascular ultrasound analysis traces arterial wall changes in abdominal aortic aneurysm

After AAA induction in *ApoE*^-/-^ mice by Ang II infusion for 28 days, abdominal aorta was assessed by HR vascular ultrasound. Morphological and functional analyses of the arterial wall were performed by VevoVasc software as described above **(Fig. 3C, D)**. Ang II infused *ApoE*^-/-^ mice showed significantly larger maximum diameters compared to controls, with distinguishable aneurysmatic segments of the abdominal aorta in HR ultrasound analysis **(Fig. 3A, B, E)**. IMT revealed to be enlarged in AAA compared to controls (101.2 ±5.9 μm vs. 136.7 ±7.1 μm; **Fig. 3F**). Moreover, PWV was increased in the aneurysmatic aorta, indicating higher arterial wall stiffness compared to control animals (1.10 ±0.11 m/s vs. 1.61 ±0.12 m/s; **Fig. 3G**). Elastic properties of the aortic wall were decreased in AAA, which presented a reduced distensibility (130.6 ±12.8 MPa^-1^ vs. 76.2 ±13.1 MPa^-1^) and decreased GRS (23.8 ±2.8% vs. 12.5 ±2.5%; **Fig. 3H, I**)

**Figure 3:**
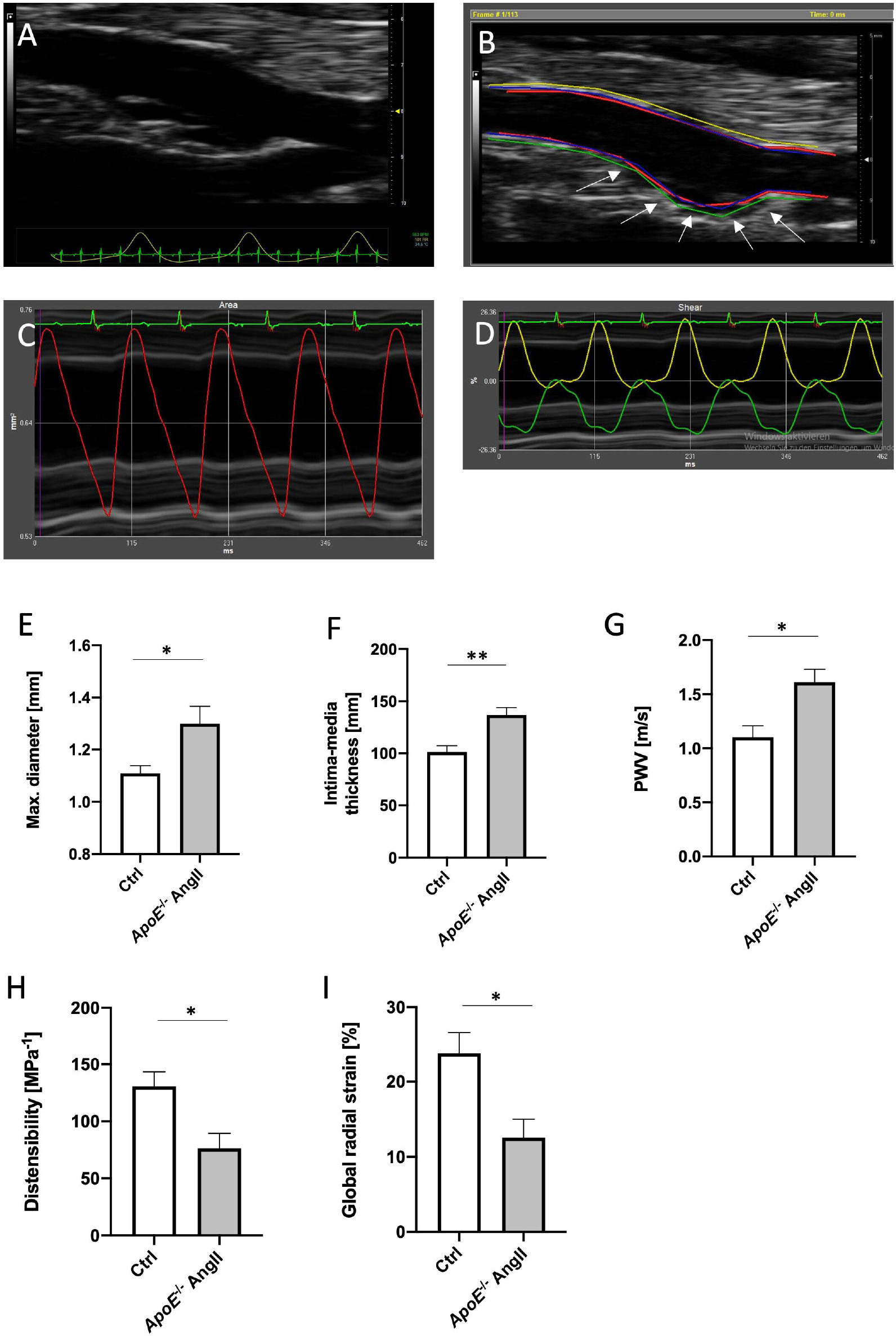
Sonographic assessment of vessel morphology and vascular function in abdominal aortic aneurysm. Abdominal aorta was displayed in B-Mode from paravertebral in control (A) and *ApoE*^-/-^ mice after 28 days of Ang II infusion (B; arrows indicate abdominal aortic aneurysm). Longitudinal images of the aorta were acquired. Borders of the vessel wall were defined in VevoVasc software and movement of lumen (C), ventral (green) and dorsal (yellow) outer vessel wall were traced by speckle tracking (D). Maximum diameter was significantly increased in *ApoE*^-/-^ mice after 28 days of Ang II infusion (E). IMT and PWV were higher in aneurysmatic diseased aortas compared to controls (F, G). Wall elasticity was significantly decreased, with curbed wall distensibility and global radial strain in Ang II treated *ApoE*^-/-^ mice (H, I). *n*=5-7; *p<0.05; **p<0.01.

### Ultrasound analysis results are consistent with histological evaluation of vascular disease models

Vessel morphology was assessed by PSR staining and collagen content was quantified as described before **(Fig. 4A)**. In the atherosclerosis model, sections of the thoracic aorta were analyzed. In line with IMT thickening in vascular ultrasound of western diet fed *ApoE*^-/-^ mice, area of the media was numerically larger in *ApoE*^-/-^ mice compared to controls, yet not reaching statistical significance **(Fig. 4B)**. However, media area correlated significantly with arterial wall distensibility and was also strongly associated with GRS as shown by simple linear regression **(Fig. 4E, F)**. Collagen content was elevated in thoracic aortas of *ApoE*^-/-^ mice, being more than twice as high as in aortic sections of control animals (38.4 ±7.7% vs. 91.7 ±1.6%; **Fig. 4C**). Simple regression analysis revealed a significant correlation of media collagen content with PWV and GRS **(Fig 4H, suppl. Fig. S1D)**. Moreover, high collagen content was associated with an increase in IMT, yet not reaching statistical significance **(Fig. 4G)**. Adventitia displayed a significant reduction in collagen fibers in *ApoE*^-/-^ mice after 12 weeks of western diet **(Fig. 4D)**. Likewise, HR vascular ultrasound analysis was compared to PSR stainings of AAA and control mice **(Fig. 5A-C)**. Area of tunica media was significantly increased in aneurysmatic abdominal aortas of *ApoE*^-/-^ mice after Ang II infusion (12.3 ±3.3 mm^2^ vs. 23.3 ±6.3 mm^2^; **Fig. 5D**) and correlated significantly with markers of arterial stiffening and IMT as measured by HR ultrasound **(Fig. 5G, H, suppl. Fig. S2A)**. Collagen content rose markedly in the media of AAA mice as indicated by automated quantification and correlated significantly with IMT measured by vascular ultrasound as assessed *via* simple regression analysis (37.2 ±6.1% vs. 71.6 ±9.5%; **Fig. 5E, I)**. High collagen content of the tunica media in trend was associated with lower GRS in AAA **(Fig. 5J)**. Again, collagen content of the adventitia was significantly lower in the aortic wall of AAA compared to controls (62.8 ±6.1% vs. 28.4 ±9.5%; **Fig. 5D**).

**Figure 4:**
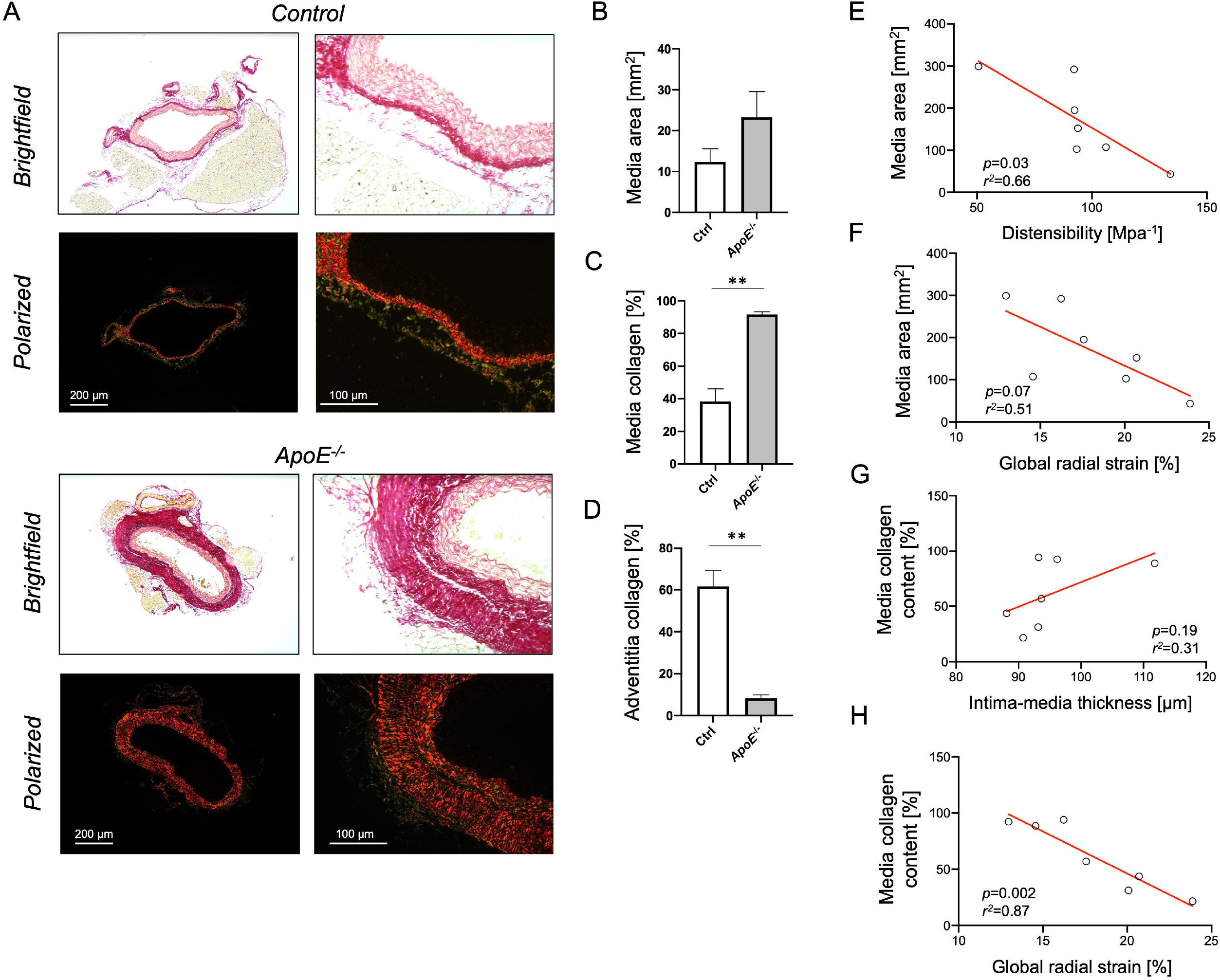
Collagen quantification in thoracic aorta after induction of atherosclerosis. Aortic sections of *ApoE*^-/-^ mice fed a western diet and controls were stained with Picro Sirius Red and collagen content was visualized in polarized light (A). Representative images are shown. Tunica media area was numerically larger and collagen content was significantly increased in atherosclerotic mice (B, C). Collagen content in adventitia was significantly lower in western diet fed *ApoE*^-/-^ mice compared to controls (D). Correlation of media area with distensibility (E) or global radial strain (F) and of media collagen content with intima-media thickness (G) and global radial strain (H) as assessed by high resolution ultrasound. Scale bars: 200μm (left panel) and 100μm (right panel) *n*=3-4 for PSR staining; **p<0.01.

**Figure 5:**
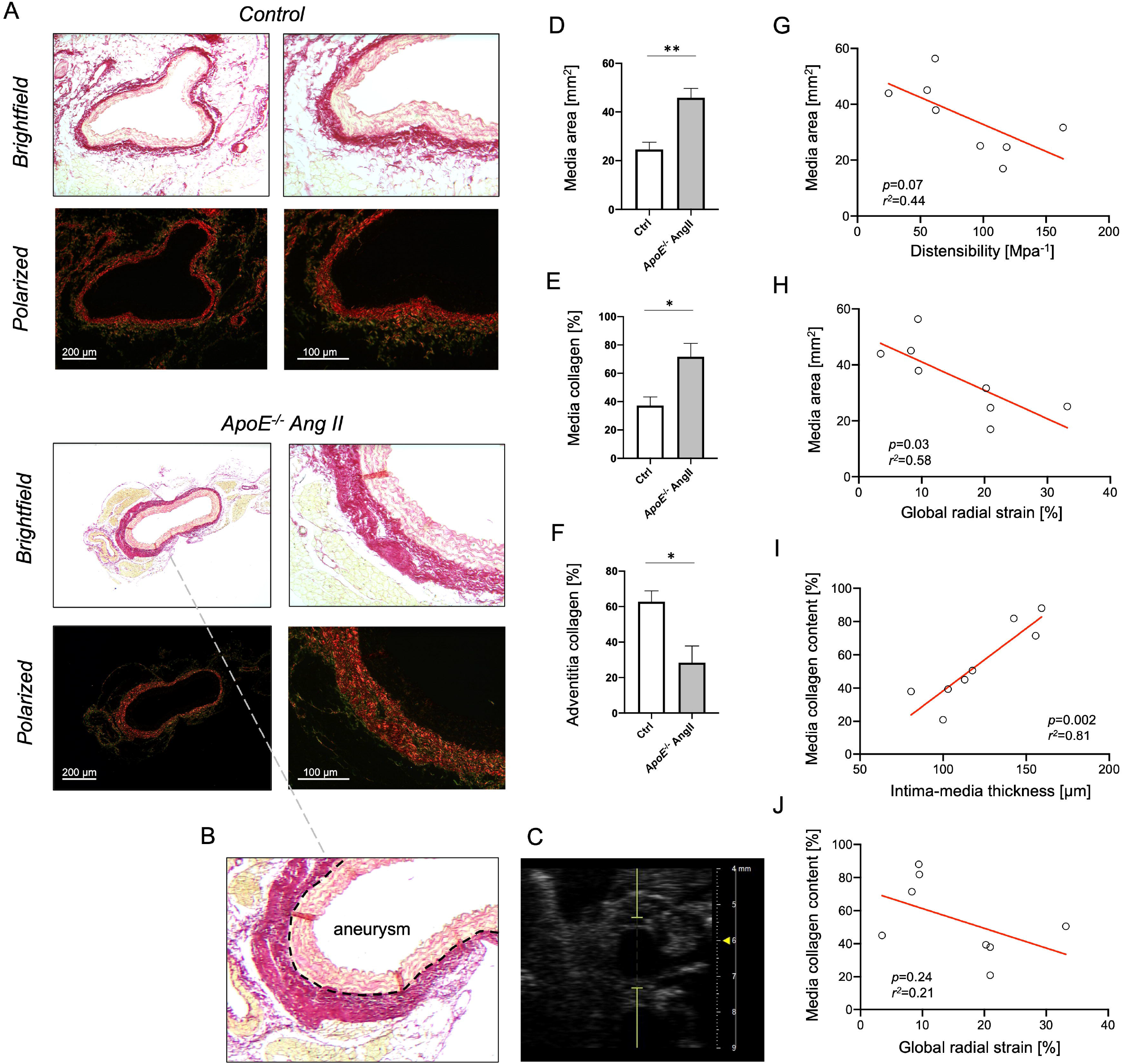
Collagen quantification in abdominal aorta in the angiotensin II aneurysm model. Representative images of control and AAA mice in brightfield and polarized light (A) with histological (B) and sonographic (C) cross section of AAA region. Picro Sirius Red staining revealed area of tunica media to be thicker in the AAA region compared to controls (D). Collagen content of the media was significantly increased in AAA (E), but lower in the tunica adventitia (F). Correlation of media area with distensibility (G) or global radial strain (H) and of media collagen content with intima-media thickness (I) and global radial strain (J) as assessed by high resolution ultrasound. Scale bars: 200μm (left panel) and 100μm (right panel) *n*=4; *p<0.05; **p<0.01.

## Discussion

In this study we show that HR vascular ultrasound 1) is a useful tool for morphologic and functional assessment of different vascular pathologies in small animal models, 2) delivers information on physiological parameters of arterial function beyond histological or *ex vivo* examinations and that 3) the obtained results match with histologic findings of aortic sections.

We here investigated mouse models of atherosclerosis and of AAA on an *ApoE*^-/-^ background. As expected, IMT of the carotid artery was broadened in *ApoE*^-/-^ mice after 12 weeks of western diet, indicating atherosclerotic wall changes [10]. Accordingly, area of the tunica media was numerically increased in the thoracic aorta of those animals compared to controls as examined by histological analyses. Further, automated collagen quantification revealed higher proportions of collagen content of the media in atherosclerotic mice. These molecular changes have been previously described as characteristic features in atherosclerosis contributing to arterial stenosis [11]. Hence, excessive collagen accumulation in media layer explains not only increased PWV as surrogate for arterial stiffness, but also reduced elasticity markers of the arterial wall as observed for distensibility and global radial strain analysis in HR vascular ultrasound. Human studies revealed arterial strain values to be reliable predictors of early atherosclerotic wall changes, significantly correlating with IMT [12]. Using latest vascular ultrasound techniques, determination of strain and elasticity parameters becomes feasible in mouse models as well, providing reliable results in terms of atherosclerotic phenotype assessment. In our study, simple regression analysis confirmed a strong correlation of different functional parameters and morphological characteristics as assessed by histological examination.

Similarly, strain analysis of the aortic wall by speckle tracking has been suggested for characterization and risk stratification of AAA in humans. Derwich *et al*., showed circumferential wall strain in AAA patients to be drastically reduced compared to young, healthy individuals [13]. With HR vascular ultrasound, we observed the same effect in *ApoE*^-/-^ mice treated with Ang II for AAA induction. Matching those findings, not only aortic wall distensibility was diminished, but also PWV was increased, suggesting a reduction in elastic properties and stiffening of the aneurysmatic aorta. Both have been described for AAA in humans and depict physiological relevance of increased collagen accumulation in the utilized AAA mouse model as observed in PSR staining and automated collagen quantification [14, 15]. However, several reports suggest aortic wall distensibility to predict the risk of infrarenal AAA rupture in later stages of disease, with higher distensibility measures correlating with a greater risk of rupture [16, 17]. Since AAA rupture also occurs in various animal models and aortic dissection or fatal rupture can be triggered, for instance by Ang II in osteoprotegerin-deficient mice, HR ultrasound might be useful to detect early aortic wall changes and predictors of aortic rupture or dissection in those models [18]. In this context, recent data suggest parameters of aortic wall stress and morphology to be even more accurate in predicting rupture risk than plain measurements of AAA diameters [19]. We found PWV and media thickness to be increased in AAA induced by Ang II in *ApoE*^-/-^ mice. Similar observations were made by Busch *et al*., describing thickness of intima-media to be elevated in surgical, elastase-induced AAA mouse models, but also in human aortas of AAA patients [20]. Likewise, PWV is increased in diseased aortas of patients with AAA [21]. HR vascular ultrasound facilitates similar risk calculations in various animal models of AAA based on the assessment of biomechanical parameters. In AAA animals, we observed a reduced collagen content of the adventitia compared to controls. Fitting those results, previous studies described collagen fibers to be disarrayed and sparse in adventitia of human AAA as assessed by electron microscopy [22], which was shown here for the *ApoE*^*-/-*^ Ang II mouse model as well.

This study has some limitations to be mentioned. First, causes of AAA development are heterogenous and reach from genetic disorders like Marfan’s disease to infectious and inflammatory conditions. Hemodynamic characteristics and changes in wall stress or morphology of the aneurysmatic aorta might differ in those diseases from the here applied *ApoE*^-/-^ Ang II mouse model, mainly inducing AAA by increased blood pressure levels and atherosclerotic predisposition. Likewise changes in collagen content and aortic stiffness might be altered in enzymatically induced AAA models and also in other models of atherosclerosis. Nonetheless, divergence of hemodynamic or morphologic features in other disease models do not seem likely. Finally, the study is limited by the small sample size and findings from animal models regarding hemodynamics and morphologic characteristics of the diseased aorta cannot be fully extrapolated to the AAA in humans. However, we here present a valid method for *in vivo* assessment of murine vascular pathologies that is consistent with established disease markers assessed by histological analysis.

## Conclusions

HR vascular ultrasound allows an accurate determination of changes in arterial wall morphology, but also elastic properties and arterial stiffness in mouse models of different vascular pathologies. We demonstrated thickening of the IMT, heightened stiffness of the arterial wall combined with reduced GRS and an impaired distensibility in an *ApoE*^*-/-*^-based atherosclerosis model and in diseased aortic segments of AAA mice. These results highlight a comprehensive elaboration and characterization of functional parameters in vascular pathologies, which were supported by automated collagen quantification, indicating media thickening with increased collagen deposition as a structural correlate of our physiological findings.

## List of Abbreviations

AAA: Abdominal aortic aneurysm
Ang II: Angiotensin II
GRS: Global radial strain
HR: high resolution
IMT: Intima-media thickness
PWV: Pulse wave velocity
PSR: Picro Sirius Red

## Author Contributions

AH, DM and HG contributed to conception and design of the study. MA, ML, JS, JN and MS performed experiments and data analysis. AH and SG performed the statistical analysis. AH wrote the first draft of the manuscript. MM, HG, DM, MA and HW wrote sections of the manuscript. All authors critically revised the manuscript, read and approved the submitted version.

## Acknowledgements

We cordially thank Mr. Max Becker for his support in performing vascular ultrasound examinations.

