## Supplement for "Vascular ultrasound for *in vivo* assessment of arterial pathologies in a murine model of atherosclerosis and aortic aneurysm"

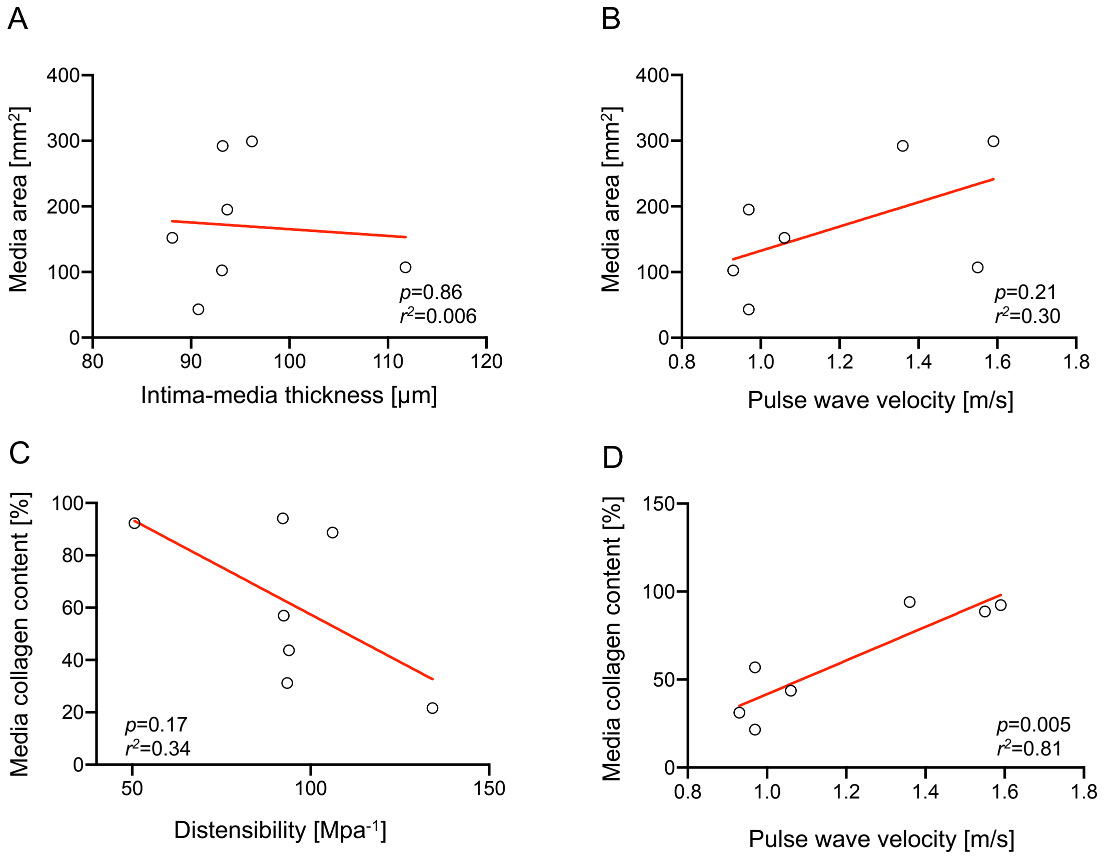


**Suppl. Figure S1: Simple regression analysis of sonographic and histological parameters in atherosclerosis.** Correlation of media area with intima-media thickness (A) and pulse wave velocity (B) and of media collagen content with distensibility (C) and pulse wave velocity (D) as measured by high resolution ultrasound. *n*=7.

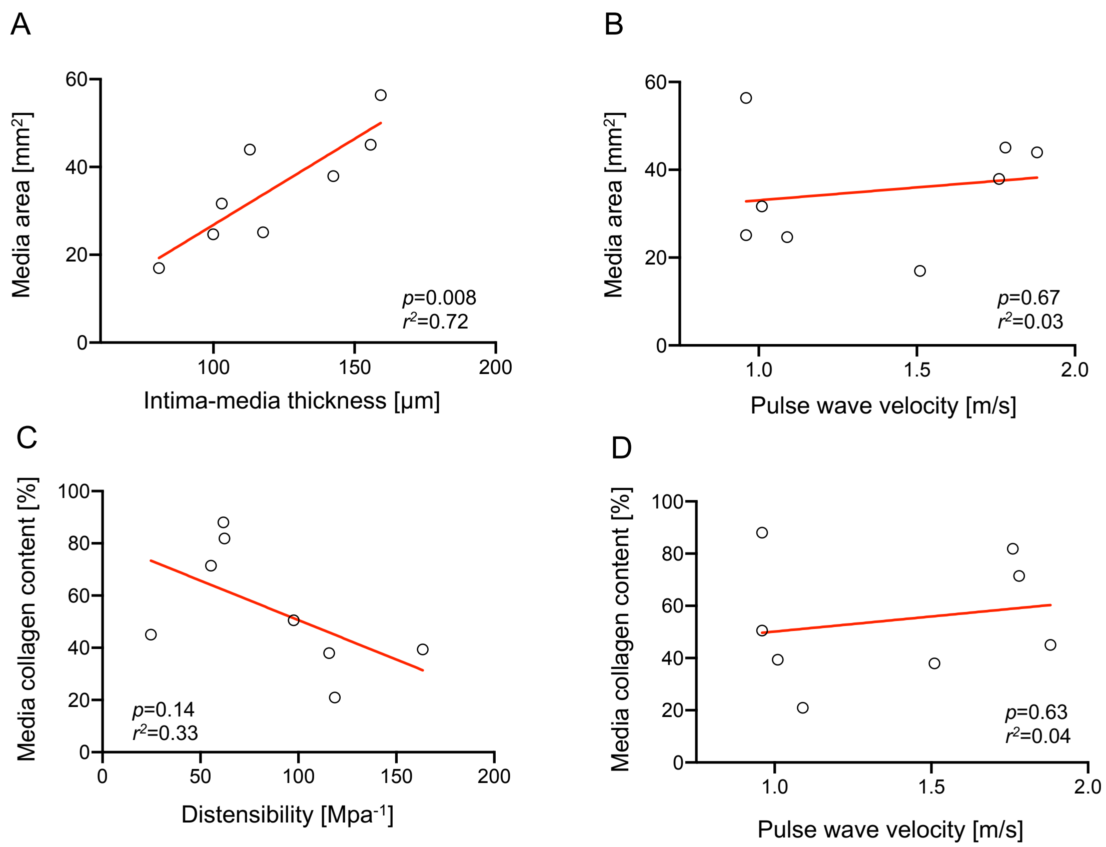


**Suppl. Figure S2: Simple regression analysis of sonographic and histological parameters in abdominal aortic aneurysm.** Correlation of media area with intima-media thickness (A) and pulse wave velocity (B) and of media collagen content with distensibility (C) and pulse wave velocity (D) as measured by high resolution ultrasound. *n*=8.
